# Prediction of migratory routes of the invasive fall armyworm in eastern China using a trajectory analytical approach

**DOI:** 10.1101/625632

**Authors:** Xi-Jie Li, Ming-Fei Wu, Jian Ma, Bo-Ya Gao, Qiu-Lin Wu, Ai-Dong Chen, Jie Liu, Yu-Ying Jiang, Bao-Ping Zhai, Regan Early, Jason W. Chapman, Gao Hu

**Author notes:** These authors have contributed equally to this work (Xi-Jie Li, Ming-Fei Wu). Correspondence to: Gao HU, College of Plant Protection, Nanjing Agricultural University, 1 Weigang Road, Nanjing 210095, China.

## Abstract

**BACKGROUND:** The fall armyworm (FAW), an invasive pest from the Americas, is rapidly spreading through the Old World, and has recently invaded the Indochinese Peninsula and southern China. In the Americas, FAW migrates from winter-breeding areas in the south into summer-breeding areas throughout North America where it is a major pest of corn. Asian populations are also likely to evolve migrations into the corn-producing regions of eastern China, where they will pose a serious threat to food security.

**RESULTS:** To evaluate the invasion risk in eastern China, the rate of expansion and future migratory range was modelled by a trajectory simulation approach, combined with flight behaviour and meteorological data. Our results predict that FAW will migrate from its new year-round breeding regions into the two main corn-producing regions of eastern China (the North China and Northeast China Plains), via two pathways. The western pathway originates in Myanmar and Yunnan, and FAW will take four migration steps to reach the North China Plain by July. Migration along the eastern pathway from Indochina and southern China progresses faster, with FAW reaching the North China Plain in three steps by June and reaching the Northeast China Plain in July.

**CONCLUSION:** Our results indicate that there is a high risk that FAW will invade the major corn-producing areas of eastern China via two migration pathways, and cause significant impacts to agricultural productivity. Information on migration pathways and timings can be used to inform integrated pest management strategies for this emerging pest.

## 1 INTRODUCTION

The fall armyworm (FAW), *Spodoptera frugiperda* (J. E. Smith), is a pest noctuid moth that principally attacks corn (maize) but has a wide host range. It is native to the New World, where it breeds continuously in tropical and sub-tropical regions of the Americas, but also has migratory populations that invade temperate North America every spring.^1, 2^ In January 2016 an outbreak of FAW was discovered in West Africa (Nigeria and Ghana), and since this initial outbreak it has spread throughout the Old World at a phenomenal rate. Within two years of arriving in West Africa it had reached almost all countries in sub-Saharan Africa.^3-5^ In May 2018, FAW were discovered in Karnataka in southwest India, and by late-2018 FAW outbreaks had been found considerably further east, in Myanmar and northern Thailand.^6-8^ Its presence in China was confirmed when larvae found in corn in southwest Yunnan province (southwest China) were identified in January 2019 as FAW.^9, 10^ By April 2019 it had spread through much of Yunnan, and also reached the southern Chinese provinces of Guangxi, Guangdong, Guizhou and Hunan (see Fig. 1), as well as Laos and Vietnam.^11, 12^

**Figure 1.**
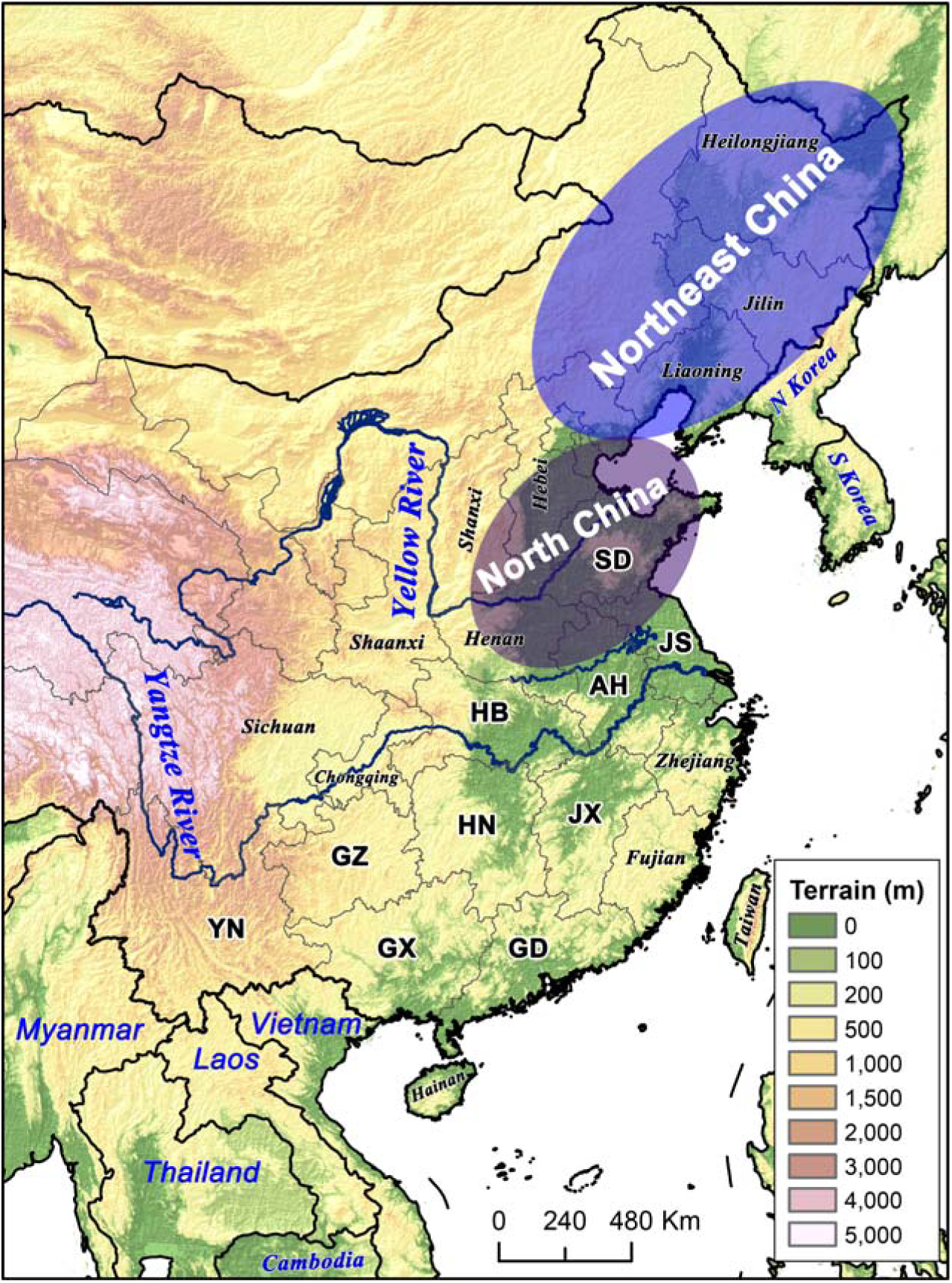
Topography of the East Asian study area. Most of eastern China is a large area of relatively flat land with few natural barriers to insect migration, but Southwest China (Yunnan and Sichuan) is a largely mountainous area with many barriers to migration. Corn is planted in each province in China, but the major corn-growing areas are the North China Plain and the Northeast China Plain. Simulated migration trajectories of FAW were started from Myanmar, Thailand, Laos, Vietnam and provinces in southwest, southeast and east-central China indicated by a 2-letter code (YN: Yunnan, GX: Guangxi, GD: Guangdong; GZ: Guizhou, HN: Hunan, JX: Jiangxi, HB: Hubei, AH: Anhui, JS: Jiangsu, and SD: Shandong). Other provinces and countries mentioned in the text are indicated on the map. The western migratory pathway originates in Myanmar and Yunnan, and passes through Guizhou, Chongqing, Sichuan and Shaanxi before merging with the eastern pathway. The eastern migratory pathway originates in northern Thailand, Laos, Vietnam, Guangxi and Guangdong, and passes through all south-eastern and east-central provinces before ultimately reaching the North China and Northeast China Plains.

FAW can survive over-winter throughout most of Southeast Asia (Myanmar, Thailand, Laos, Cambodia and Vietnam) and also in the sub-tropical provinces of China (Yunnan, Guangxi, Guangdong, Hainan, Fujian and Taiwan) lying approximately south of the Tropic of Cancer.^13^ It is highly likely that FAW populations breeding year-round in these regions will evolve annual spring migrations northwards into eastern China (and presumably south again the following autumn), just as FAW populations in North America migrate annually between the northernmost winter-breeding areas (south Texas and south Florida) and the northern United States. ^1, 2^

The caterpillars of FAW have a very wide host range, and are known to damage more than 180 species of plants.^14^ Corn is the preferred host, and yield losses of between 15–73% are typically caused by FAW outbreaks in corn.^14, 15^ Recent studies of projected yield loss in Africa, combined across twelve major corn-producing sub-Saharan countries, indicated that between 4.1–17.7 million tons of corn, with a value of $1.09–4.66 billion, will be lost annually due to the newly-invasive FAW populations.^3, 4^ China is the second largest corn producer in the world, and corn is the third commonest crop after rice and wheat in China, where it is grown in all provinces. The main corn-growing areas are the North China Plain (mainly the provinces of Henan, Shandong and Hebei, see Fig. 1) and the Northeast China Plain (Liaoning, Jilin and Heilongjiang, see Fig. 1) in eastern China, and these regions (plus the Korean Peninsula and Japan) are potentially suitable for summer-breeding populations^13^ if FAW can reach them. Therefore, Chinese agricultural production and food security will be seriously threatened if FAW evolve a regular migratory route which will allow them to exploit the principal corn-producing regions of East Asia to the north and east of the current distribution.

International trade is considered to be an important cause of the rapid expansion of FAW.^13^ In addition, this species has the capability to achieve natural long-distance range expansion, as adults can migrate hundreds or even thousands of kilometres on high-altitude winds over several successive nights;^1, 2^ for example, FAW were reported to be transported by low-level jets from Mississippi in the southern United States to southern Canada, a distance of 1,600 kilometers.^16^ Although it is unlikely that natural windborne migration was responsible for the moths crossing the Atlantic and Indian Oceans to colonize Africa and India respectively, natural migration is hugely important for their spread within Africa, and during their invasion of East and Southeast Asia.^4^ European countries are worried about the very real possibility that the moths will migrate to Europe after they breed successfully in North Africa.^17^

Now that FAW have arrived in Southeast Asia and southern China, there is a very high possibility that they will invade eastern China on an annual basis. Two main migratory routes are possible: a western and an eastern route. The western route involves windborne transport from the westerly winter-breeding region (Myanmar / Yunnan), via Guizhou and Sichuan and on into eastern China (Fig. 1). The eastern route originates from the easterly winter-breeding region (northern Thailand, Laos, Vietnam, Guangxi and Guangdong), and involves transport on favourable winds associated with movement of the Asian monsoon via east-central China, and on into the main corn producing areas (the North China and Northeast China Plains) (Fig. 1). The eastern route is the important migratory pathway for many migratory pest moths in China, including beet armyworm *Spodoptera exigua*,^18^ beet webworm *Loxostege sticticalis*,^19^ cotton bollworm *Helicoverpa armigera*,^20^ Oriental armyworm *Mythimna separata*^21-24^ and rice leaf roller *Cnaphalocrocis medinalis*.^25^ As is the case for these other migratory pests, at these latitudes FAW can only breed successfully in the summer and cannot survive overwinter, and so these regions will need to be reinvaded on an annual basis.^2, 13, 26^ Hence, the question of whether FAW can evolve a regular, seasonal round-trip migration between the year-round breeding zone in Southeast Asia / southern China, and the potential summer-breeding zones in North and Northeast China is the key to whether they can cause frequent and wide-scale crop damage in China. However, East Asia would appear to be a very suitable region for the development of long-distance annual migrations of FAW, for four reasons. Firstly, the maize producing regions of eastern China lie at a similar latitude and have similar climate to the FAW native migratory range in the USA. Secondly, East Asia has a wide extent of tropical and subtropical regions on the Indochina Peninsula and in southern China, which provide a favourable environment for FAW to maintain large populations over the winter. Thirdly, there is a continuous agricultural ecosystem spanning a large latitude range in Southeast and East Asia with year-round production of suitable crops (corn, sugarcane, rice, etc) enabling continuous breeding if FAW can move between regions. Finally, the annual East Asian summer monsoon provides a ‘highway’ of favourable winds for the airborne transport of migratory organisms, towards the north in the spring and returning south in the autumn. Taken together, this means the recent colonisation of Southeast Asia and southern China is very likely to result in the emergence of a round-trip migratory cycle that will exploit the seasonal resources available in eastern China. China is therefore facing a great risk to its food security and agricultural productivity due to the invasion of FAW into the region. It is thus important to identify the migration routes, timing of the seasonal movements, and potential summer-breeding range of FAW in eastern China, in order to design strategies to monitor and control this pest. In this study we predict the future migratory pathways of FAW using trajectory simulations modified to take account of FAW migration behaviour.

## 2 METHODS

We identified the potential endpoints of FAW moth migrations by calculating forward flight trajectories from source areas where FAW are currently known to be breeding, or from potential future source areas we predict they will breed in the near future. To improve the accuracy of the trajectory simulations, we developed a new numerical trajectory model that takes account of flight behaviour and self-powered flight vectors (as these are known to substantially alter trajectory pathway^27, 28^), and trajectory calculation is driven by high spatio-temporal resolution weather conditions simulated by the Weather Research and Forecasting (WRF) model.^29^ This trajectory model has been used successfully for many other insect migrants, such as corn earworm (*Helicoverpa zea*), Oriental armyworm, rice leaf roller, and rice planthoppers.^24, 25, 29-33^ The program for calculating trajectories was designed in FORTRAN ^24, 29, 31^ and run under CentOS 7.4 on a server platform (IBM system x3500 M4).

### 2.1. Weather Research and Forecasting model

The Weather Research and Forecasting (WRF) model (version 3.8, www.wrf-model.org) was used to produce a high-resolution atmospheric background for trajectory calculation. The WRF is an advanced meso-scale numerical weather prediction system (https://www.mmm.ucar.edu/weather-research-and-forecasting-model).^34^ In this study, the dimensions of the model domain were 140 ×150 grid points at a resolution of 30 km. Twenty-nine vertical layers were available and the model ceiling was 100 hPa. More detail of the scheme selection and parameters for the modelling are listed in Supplementary Table S1 and Fig. S1. National Centers for Environmental Prediction (NCEP) Final Analysis (FNL) data was used as the meteorological data for the model input. FNL is a six-hourly, global, 1-degree grid meteorological dataset. The model forecast time is 72 h with data outputs at 1 h intervals, for horizontal and vertical wind speeds, temperature and precipitation.

### 2.2. Self-powered flight behaviours of FAW

The flight behaviour of FAW were included in the trajectory simulation by making the following assumptions. (i) Nocturnal moths perform ‘multi-stop’ migration, in which moths only take off at dusk, terminate migratory flight the following dawn, and then take-off again at the next dusk.^25, 27, 28^ FAW here was assumed to take off at 20:00 Beijing Time (BJT), stop at 06:00 BJT, and fly for three consecutive nights whenever temperature conditions were suitable (see below). (ii) Other species of similar-sized noctuid moth pests have a self-powered flight speed of about 2.5–4 m/s.^27, 35, 36^ Therefore, we added a self-powered flight vector of 3.0 m/s in the trajectory modelling. As we don’t know if the Asian FAW moths have a preferred flight heading, we assumed that the flight vector will be aligned with the downwind direction. (iii) Radar studies of FAW in the USA ^1, 37-39^, and of similar noctuid moth pests elsewhere ^27, 35^, show that these moths typically migrate at the altitude of the of the low-level jet where wind speeds are relatively fast (often >10 m/s). We did not explore altitudinal profiles of wind speeds before trajectory modelling, and thus to ensure we would capture the most likely flight height, we started trajectories from eight different altitudes: 500, 750, 1000, 1250, 1500, 1750, 2000 and 2250 m above mean sea level (amsl). In the eastern pathway we only calculated trajectories at heights from 500–1500 m amsl as ground heights in this region are relatively low, but we used all 8 altitudes for the western pathway as much of the land in this region (particularly in Yunnan) is >1000 m amsl. We assumed that FAW cannot fly when the air temperature at flight altitude falls below 13.8 °C, the minimum temperature for survival of FAW ^13, 40^, and so trajectories were terminated on any night/height combination which dropped below this temperature.

### 2.3. Departure points for forward trajectories

We investigated the two main potential migratory pathways (the western and eastern routes) by which FAW may annually invade eastern China, during four separate waves of migration (March–April, April–May, May–June, and July). The western route originates in Myanmar and Yunnan, and develops via Guizhou and Sichuan (Fig. 1). To model this route, trajectories were started from all potential departure points at every 1° grid for the following schemes: from (i) Myanmar and Yunnan in March–April; (ii) Yunnan in May; (iii) Yunnan and Guizhou in June; and (iv) Yunnan and Guizhou in July (Fig. 2, Fig. S1). Myanmar and Yunnan were selected due to the fact that FAW has been present during the first winter period of 2019, and Guizhou was selected because many trajectories from Yunnan reached this province in May.

**Figure 2.**
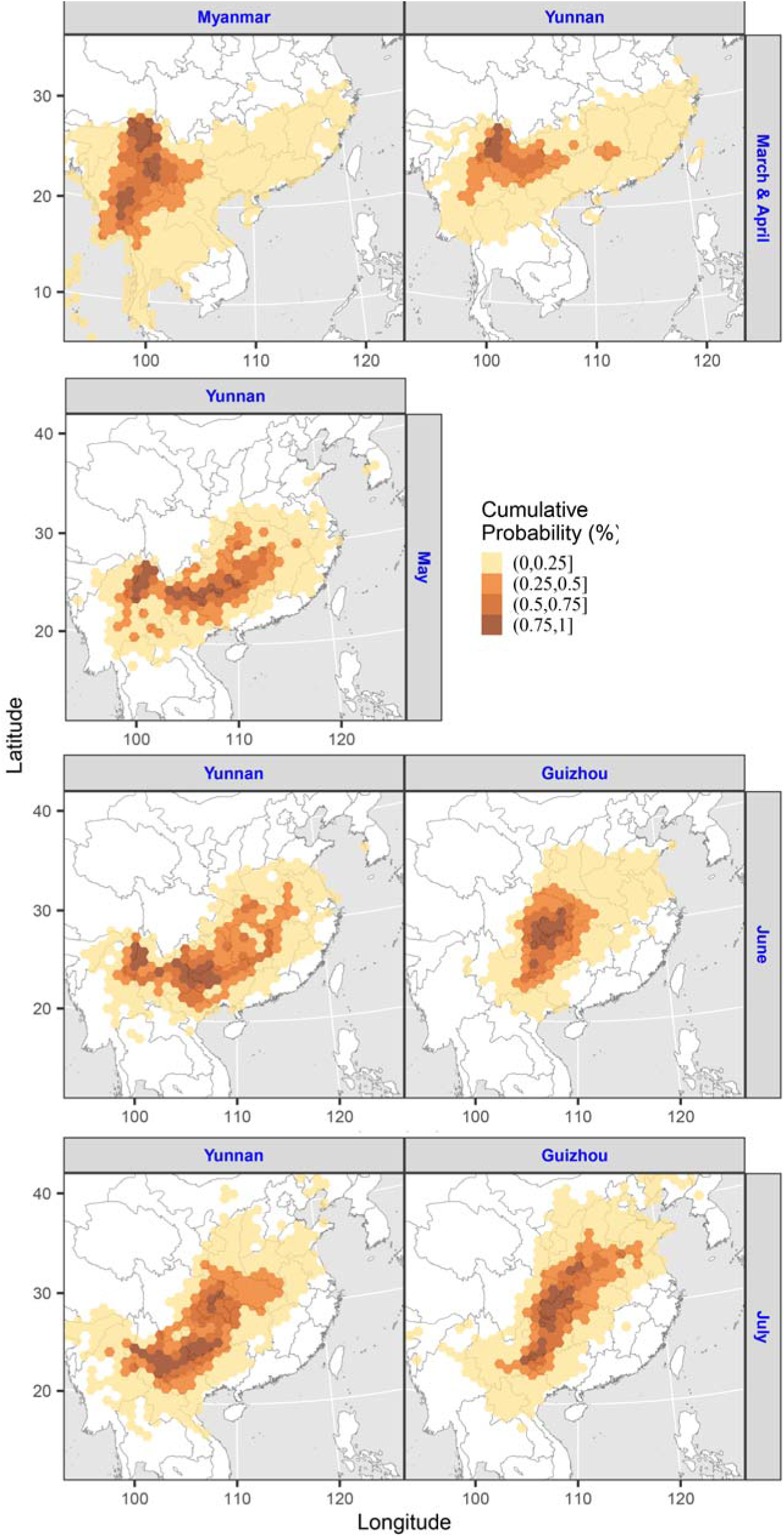
Distribution of endpoints of FAW forward migration trajectories along the western migratory pathway. The start-points and time periods of trajectories are labelled on the top / right of each panel. Trajectory analyses were conducted over three consecutive nights, and only the final endpoint of each 3-nights trajectory is shown. Each hexagonal cell covers 10,000 km^2^.

The eastern route starts in northern Indochina, Guangxi and Guangdong, and develops via east-central China towards the main corn-producing regions in North and Northeast China (Fig. 1). To model this route, trajectories were started from all potential departure points at every 1° grid for the following schemes: from (i) Thailand, and Laos / Vietnam, in March–April; (ii) Guangxi and Guangdong in April–May; (iii) Hunan / Jiangxi, and south Hubei / south Anhui, in May–June; and (iv) Hubei / Anhui, and Jiangsu / Shandong, in July (Fig. 2, Fig. S1). The first two schemes were selected based on current (April 2019) distribution of FAW, while the latter two schemes were selected based on the results of the trajectories originating from the first two schemes earlier in the season.

We simulated the FAW trajectories by using average meteorological conditions at flight altitude from the past 5 years (2014–2018). In total, >0.6 million trajectories were calculated (Table S2), making this the largest study of FAW migration pathways conducted.

### 2.4. Effect of flight altitude on migration trajectories

To investigate whether flight altitude would have affected distance and directional components of the trajectories, we carried out a comparative analysis to see how three migration parameters varied with altitude. Firstly, we calculated the average distance travelled during the three nights of migratory flight at each of the modelled flight heights (between 500 and 2250 m amsl in the western pathway, and between 500 and 1500 m amsl in the eastern pathway), to see how distance varied with height across the regions and seasons. Secondly, we looked at how the mean direction of the trajectories varied with altitude. Thirdly, we investigated the degree of directional spread of the trajectories with altitude. For each altitude, we used the Rayleigh test for circular data ^41^ to calculate the mean direction and the r-value of the circular distribution of the directions of the trajectory endpoints from the starting locations. The Rayleigh r-value ranges from 0 to 1, with higher values indicating a greater clustering of directions around the mean and lower values indicating a wider angular spread of trajectory endpoints. These three parameters therefore indicate the effect that flight altitude selection will have on (i) the distance travelled during migratory flights, (ii) the mean direction of windborne transport, and (iii) the degree of dispersion or concentration that will occur over many nights of migratory flight.

## 3. RESULTS

### 3.1. The Western Migratory Pathway

The first detection of FAW in the East/Southeast Asian region occurred in Myanmar and Yunnan (in the winter period of 2018/19) ^9, 10^, so we ran our first trajectories from these areas during March–April. In both cases, the endpoints of these trajectories (the first wave of migration) largely remained within Yunnan province indicating a rather slow northward spread (Fig. 2). However, interestingly, some trajectories from Yunnan reached the southeast corner of Guizhou province in this period, and this coincided precisely with the location of a FAW outbreak discovered in late-April 2019.^12^ The second wave of migration moved much further from Yunnan, with many trajectories ending in Guizhou (Fig. 2) and yet others travelling further east where they entered the eastern migratory pathway (see below). During June (the third wave of migration), trajectories from Guizhou moved in a northwards direction and FAW arrived in central China (eastern Sichuan, Chongqing and southern Shaanxi). The fourth wave of migration during July took FAW into the more easterly provinces of southern Shanxi, Henan and southern Shandong (Fig. 2). Our trajectory simulations therefore show that FAW moths migrating along the western pathway will reach the North China Plain during the fourth wave of migration (by July).

### 3.2. The Eastern Migratory Pathway

Migration trajectories originating from Thailand, and from Laos / Vietnam, during March–April (the first migration wave) had a high probability of ending in southern China. Trajectories from Thailand reaching China were concentrated mostly in Guangxi, while those from Laos / Vietnam also had many endpoints in Guangxi, but in addition extended further north and east, into most of Guangdong and also the southern parts of Hunan and Jiangxi (Fig. 3). During the next stage of trajectories (the second migration wave), modelled from Guangxi and Guangdong during April–May, FAW were predicted to continue travelling further north and east into China, reaching the southern fringe of the Yangtze River Valley. Guangxi trajectories were directed to the northeast and terminated mainly in Hunan, but with many endpoints also in Jiangxi and the southern regions of Hubei and Anhui (Fig. 3). Trajectories from Guangdong had a more easterly component, and were concentrated in Jiangxi, Fujian and the southern part of Zhejiang (Fig. 3).

**Figure 3.**
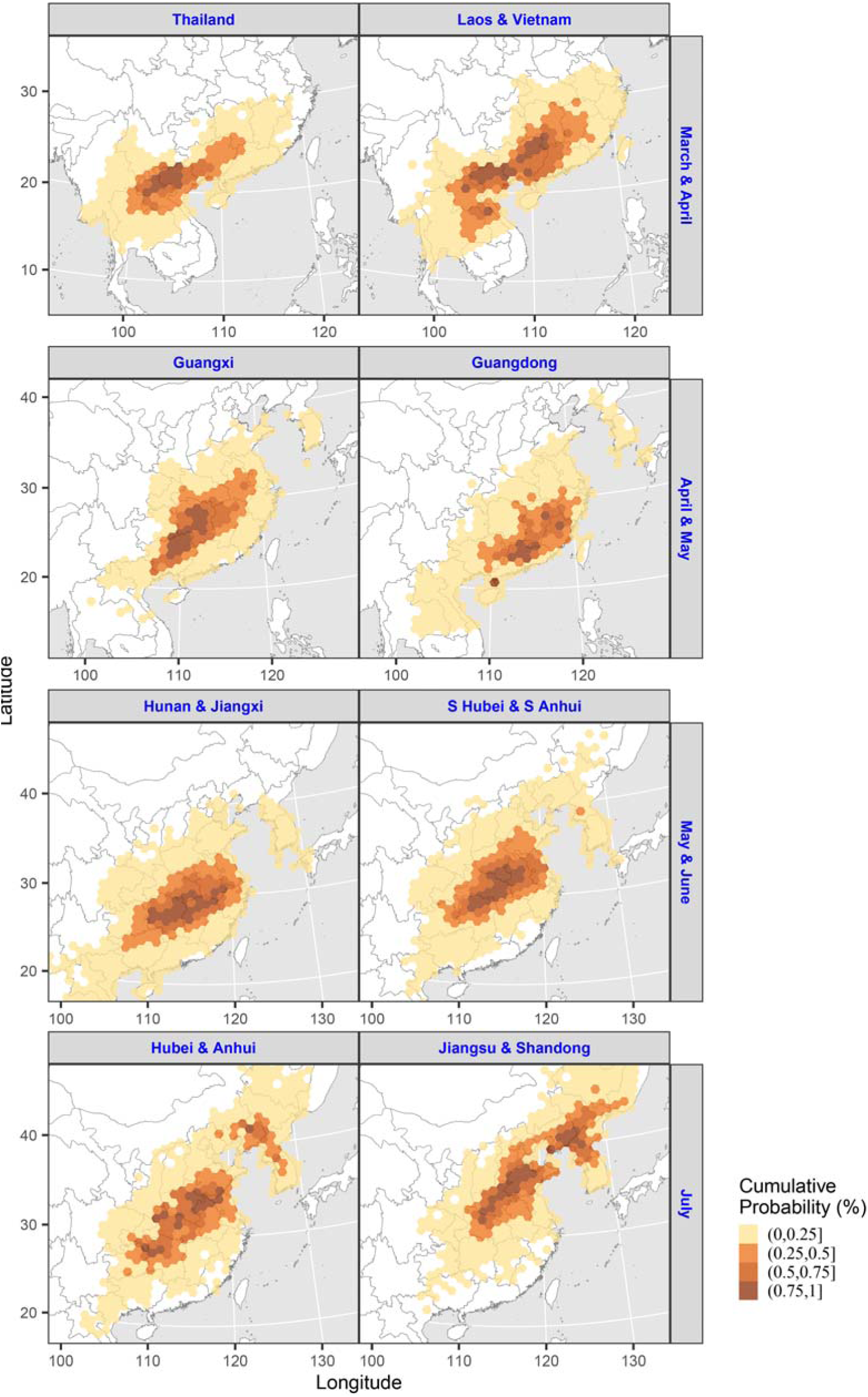
Distribution of endpoints of FAW forward migration trajectories along the eastern migratory pathway. The start-points and time periods of trajectories are labelled on the top / right of each panel. Trajectory analyses were conducted over three consecutive nights, and only the final endpoint of each 3-nights trajectory is shown. Each hexagonal cell covers 10,000 km^2^.

The third wave of migration was modelled from the Hunan / Jiangxi region, and the south Hubei / south Anhui region, during May–June. The northward progression of the migration continued in this period, although the distance travelled was relatively small and trajectory endpoints were mostly concentrated in the region between the Yangtze and Yellow River Valleys, in the provinces of Hubei, Anhui, Henan, Jiangsu and Shandong (Fig. 3). This partly overlaps with the important corn-growing region of the North China Plain (Fig. 1). The fourth wave of migration during July involved a longer distance movement to the northeast than in the third wave. Trajectories originating in Hubei / Anhui, and in Jiangsu / Shandong, extended to the northern part of the North China Plain (Hebei), and also reached important corn-growing regions in the Northeast China plain (Liaoning and Jilin) and North Korea (Fig. 3). Our trajectory simulations therefore show that FAW moths migrating along the eastern pathway will reach the North China Plain during the third wave of migration (by June, that is a month earlier than the western pathway), and will then reach the Northeast China Plain during the fourth wave (in July).

### 3.2. Effect of flight altitude on migration trajectories

In order to assess the role that flight altitude selection may have on migration pathways, we analysed how distance, direction and degree of directional clustering of the trajectories varied with altitude at each location (Fig. 4, Table S3). Trajectory height had a strong effect on the distance travelled at some locations, but the direction of the trend with altitude varied between sites, and in other regions there was no effect of altitude. In the western flyway, early in the season most trajectories from Myanmar and Yunnan were comparatively short irrespective of flight height (Fig. 4, Table S3) due to relatively cool air temperatures, which explains why the initial northward spread from this region was rather slow during March–April (Fig. 2). Later in the season however, as air temperatures warmed, flight altitude had a large effect on distance travelled, with trajectories at heights >1500 m producing considerably longer trajectories than lower altitudes in Yunnan (typically 800–1000 km versus <500 km), but with the opposite trend in Guizhou where flight below 1000 m produced the longest trajectories (Fig. 4, Table S3). Directions varied with altitude in a complicated fashion across the different regions and time periods (Table S3). The degree of directional clustering of trajectories tended to follow a regular pattern, with tighter distributions occurring at high and low altitudes, but with a much greater degree of dispersion at intermediate heights (Fig. 4).

**Figure 4.**
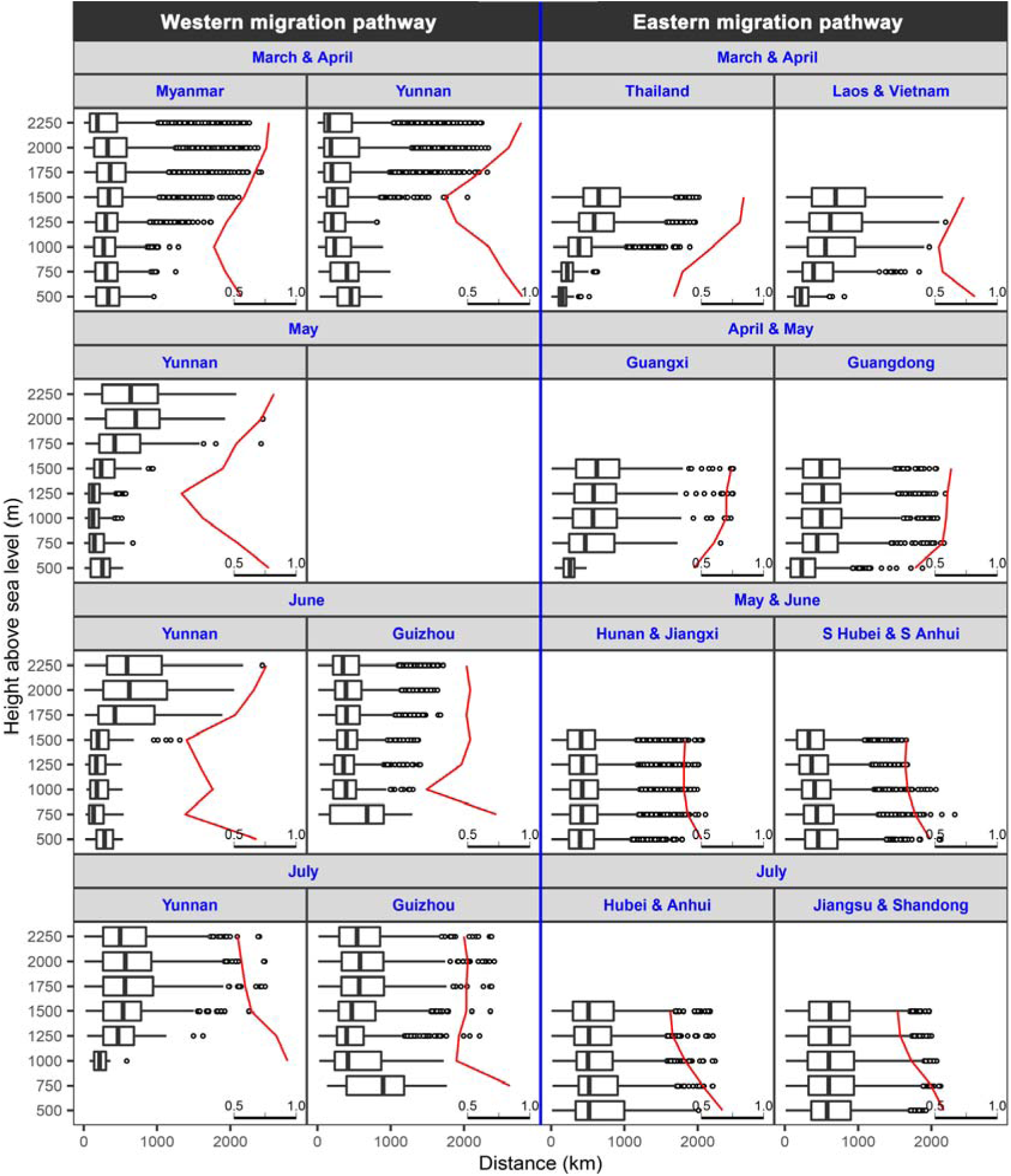
The effect of flight altitude on trajectory parameters. The black box plots show the straight-line distances between the start-points and the final endpoints for each trajectory, and how they vary with altitude. In the black box plots, central bars represent median values, boxes represent the inter-quartile range (IQR), whiskers extend to observations within ±1.5 times the IQR, and dots represent outliers. The red lines (on a secondary scale) show the Rayleigh test *r*-values for the trajectory directions at each altitude. This provides a measure of the degree of clustering of the angular distribution of directions around the mean, ranging from 0 to 1, with higher values indicating tighter clustering and thus a higher degree of common trajectory directions and lower values indicating a greater dispersion of trajectories.

Along the eastern pathway, trajectories tended to become longer and more tightly clustered with increasing altitude in the Indochina Peninsula and southern China during spring (Fig. 4). However, this patterns changed during late-spring and summer as the moths moved further north into eastern China, with trajectory distance showing no pattern with altitude but trajectory directions becoming more dispersed with increasing altitude in the Yangtze and Yellow River Valleys and the North China Plain (Fig. 4). Once again, directions varied in a complicated manner with altitude (Table S3).

## 4. DISCUSSION

In this study, we predicted future migration pathways of FAW in eastern China using a trajectory analysis approach, combined with flight behaviour of FAW and meteorological data from the past 5 years. Our results show that FAW will likely undertake annual migrations from its new overwintering area in the Indochina Peninsula and South China into the two main corn-producing areas of eastern China. The North China Plain (mainly Henan, Shandong and Hebei) is predicted to be invaded in June each year after three waves of migration along the eastern pathway, and then to receive another influx in July due to a fourth wave of migrants coming from the western pathway. The Northeast China Plain (Liaoning, Jilin and Heilongjiang) will then be invaded by a fourth wave of migrants in July that originate for the population colonising North China a month previously. This likely annual migration pathway will result in substantial damage and economic losses to corn production in these two vitally important areas unless the FAW population can be effectively managed.

Many species of insect carry out similar seasonal long-distance migrations in East Asia ^42^, including the most serious crop pests in this region, such as the oriental armyworm, beet armyworm, cotton bollworm, rice leaf roller and rice planthoppers (*Nilaparvata lugens* and *Sogatella fucifera*). Entomological radar studies have shown that the smaller, relatively weak-flying species, such as the rice leaf roller and planthoppers, do not have adaptive, wind-related, preferred flight headings or flight altitudes, and simply fly with random orientation at the altitude where they reach their flight temperature threshold.^43-45^ This means these species will be passively transported downwind, with little or no influence over their migration trajectories.^42, 46^ However, these weak-flying insects are still capable of carrying out annual round-trip migrations between their winter-breeding regions in Southeast Asia / South China, and summer-breeding regions much further north in East Asia. This is because they can benefit from the seasonally-favourable winds that dominate in this region, due to the passage of the East Asian monsoon.^30, 47^ This persistent large-scale weather system produces frequent winds from the southwest in the spring and summer, and then switches to frequent winds from the north in the autumn, over the entire East Asian migration arena, thus providing suitable transporting flows for insect migrants over the whole flight season.^43, 47^ Our study of likely FAW migration trajectories is entirely consistent with this situation, and our modelling suggests that FAW only need to take-off and climb to a few hundred meters above ground to achieve rapid, long-distance transport towards eastern China during the spring. The migration system can therefore evolve without any further specialised behaviours, simply due to the high frequency of seasonally-favourable tailwinds. Presumably the progeny of the fourth wave will start to return to the south from August onwards, though this idea still needs to be formally tested.

Simple reliance on seasonal patterns of suitable winds however is still a rather risky and inefficient strategy, and more powerful fliers (such as noctuid moths) could considerably improve the efficiency of their migratory flights, and reduce migration-related mortality ^28^, by adopting beneficial flight behaviours. Radar studies of moth migration in Europe ^35, 42, 48^ have clearly demonstrated that a closely related species of migrant moth, the silver Y *Autographa gamma*, has a syndrome of related behavioural traits which significantly increase the speed, distance, directionality and success of its migratory flights. These flight behaviours include the ability to detect and respond to the downwind direction, restricting migration to nights with seasonally-favourable high-altitude tailwinds, selecting the altitude with the fastest wind, and common orientation in seasonally-preferred migration directions.^42, 49-52^ There is growing evidence that these behaviours are probably widespread in larger insect migrants ^53, 54^, including Asian pest moths such as Oriental armyworm and cotton bollworm.^20, 21^ It would thus seem very likely that FAW populations in Asia will already have, or will rapidly evolve, some (or all) of these behaviours, and these flight behaviours will have a major impact on their trajectories.

In our trajectories the only flight behaviour we encoded into our model was a self-powered flight vector of 3 m/s in the downwind direction, whichever way the wind blew. We did not allow moths to be selective of whether to migrate or not (depending on the wind direction), nor did we allow them to orientate in seasonally-beneficial directions or select flight altitudes based on wind speed. These decisions were made simply because we know virtually nothing about the flight behaviour of the FAW populations in Asia, and we felt it safer not to make too many assumptions for the purpose of this study. However, our preliminary exploration of the impact some of these behaviours can have on migration trajectories (see Fig. 4) clearly shows that an understanding of flight behaviour will be crucial for accurately predicting the migration pathways and future range of this moth in East Asia. Behavioural studies of FAW populations in southern China should thus be carried out as a matter of urgency.

There are many similarities in the ecology and biology of FAW and Oriental armyworm, including their migratory capability, body size and self-powered flight speed, wide host range and pest status, and latitudinal extent of their breeding ranges, and thus it may be assumed that the two species will have a similar migration pattern and phenology in East Asia. The Oriental armyworm typically has only two steps in its northwards migration into northeast China. The first step involves migration from its overwintering area south of the Yangtze River into the plains between the Yangtze River and Yellow River (30°–35°N) in March and April. The next generation then migrates as far north as Northeast China and eastern Inner Mongolia, in a single step by May–June.^22, 23, 55^ However, our results indicate that FAW will require three migration steps to reach the North China Plain in June, and four steps to reach Northeast China in July. Thus its migration pattern is quite different from that of the Oriental armyworm, presumably due to differences in their minimum temperature for survival: 13.8°C for FAW, but only 9.6°C for Oriental armyworm.^13, 40, 56^ The year-round distribution of FAW in East Asia is restricted to the relatively warm and moist regions found on the Indochina Peninsula and in southern China (to the south of the Tropic of Cancer) ^13^, similar to rice planthoppers and the rice leaf roller.^25, 46^ Oriental armyworm on the other hand can survive over winter in the region south of the Yangtze River (33 °N) in China, considerably further north than FAW.^57^ Due to their similar body size (and thus flight capability and speed), and similar developmental periods (about one month per generation under suitable temperature conditions), it is expected they will achieve similar migration distances each year, and thus the occurrence area of FAW will be further south than the Oriental armyworm at any one time.

The East Asian migration arena would appear to be a highly suitable environment for the FAW, having suitable wind regimes for migration, suitable climate for rapid development and widespread availability of corn. However, other factors may influence its spread throughout the region, including distribution of alternative host plants, natural enemies and competitors. The phenotype of FAW in Africa, Myanmar and Yunnan has been identified as the corn strain, and the rice strain appears to be largely absent.^25, 46^ However, as FAW populations arrive in South China, they will encounter large areas of rice paddies, and relatively infrequent corn cultivation, which will affect its population growth. Another factor which will determine population growth is the prevalence of natural enemies, which may be expected to be low for a newly-invasive species. However, field surveys in Yunnan found that 15–20% of FAW caterpillars were infected by parasitoid wasps (unpublished data from G.P Li in Henan Academy of Agricultural Sciences), which is encouraging from the perspective of population suppression via natural biological control. In addition, FAW populations in East Asia will also encounter new competitors such as the Oriental armyworm and the Asian corn borer *Ostrinia furnacalis*. None of these factors were considered in our trajectory modelling, and we believe that ecological studies of FAW populations as they colonise East Asia should be undertaken as a priority.

In conclusion, the major corn-growing regions of China face a high risk of invasion by FAW. The North China and Northeast China Plains can be invaded by FAW via a series of 3–4 steps of northward migration, which will allow FAW to reach as far north as the border of Jilin with Heilongjiang by July. The most efficient way to prevent invasive species from entering a new country is efficient border quarantine. However, the ability of FAW to carry out long-range, windborne migration means that traditional methods of surveillance and quarantine are useless. When this study was began in January 2019, the FAW was only known from Myanmar and Yunnan, and we wanted to know if it could invade the rest of the Southeast and East Asian areas. In the intervening 4 months before this paper was submitted in May 2019, FAW had already spread to Thailand, Laos, Vietnam, Guangxi, Guangdong, Guizhou and Hunan, and its continuing spread through China to the north and east seems inevitable. Additional studies on its migration patterns, flight behaviour, ecology, and pest management are urgently required.

## ACKNOWLEDGMENTS

This work was supported though grants to G.H. by the National Natural Science Foundation of China (31822043) and the Natural Science Foundation of Jiangsu Province (BK20170026). B.G.’s visiting scholarship to the University of Exeter was funded by the China Scholarship Council and the Jiangsu Graduate Research and Innovation Projects. J.W.C. was supported by the Science and Technology Facilities Council (STFC) Newton Agritech Project “Integrating advanced earth observation and environmental information for sustainable management of crop pests and diseases” (ST/N006712/1) and the National Natural Science Foundation of China (61661136004).

