## Supplementary material for "Prediction of migratory routes of the invasive fall armyworm in eastern China using a trajectory analytical approach": Table S1, S2 and S3, Figure S3

Submission to *Pest Management Science*

**Supplementary Table S1.** Selection of scheme and parameters for the Weather Research and Forecasting (WRF) Model. Domain 1 was used in the trajectory simulation for Indochina and Yunnan in March and April, and Domain 2 was used for China from April to July.

| Item | Domain 1 | Domain 1 |
| --- | --- | --- |
| Location | 23°N, 107°E | 32°N, 108°E |
| The number of grid points | 130*150 | 140*150 |
| Distance (km) between grid points | 30 | 30 |
| Layers | 29 | 29 |
| Map projection | Lambert | Lambert |
| Microphysics scheme | WSM3 | WSM3 |
| Longwave radiation scheme | RRTM | RRTM |
| Shortwave radiation scheme | Dudhia | Dudhia |
| Surface layer scheme | Monin-Obukhov | Monin-Obukhov |
| Land/water surface scheme | Noah | Noah |
| Planetary boundary layer scheme | YSU | YSU |
| Cumulus parameterization | Kain-Fritsch (new Eta) | Kain-Fritsch (new Eta) |
| Forecast time | 72 h | 72 h |

**Supplementary Table S2.** The number of FAW forward trajectories simulated. Forward trajectories were calculated for three consecutive nights with eight different initial flight altitude heights: 500, 750, 1000, 1250, 1500, 1750, 2000 and 2250 m above mean sea level. Trajectories were terminated if: (i) low temperatures were encountered (defined as air temperatures at flight altitude below 13.8 °C); or (ii) ground height exceeds the trajectory altitude (the land frequently rises above 1000 m in Yunnan and Laos). Migration over the sea was not considered in this study, and final endpoints after 3 consecutive nights of flight that were located in the sea were deleted. In total, 624,933 trajectories were calculated, but only 186,492 (29.84%) trajectories were valid and presented in Figs. 2 and 3.

| Region | Period | Total | First night | | | Second night | | | Third Night | | | Enter sea | Normal |
| --- | --- | --- | --- | --- | --- | --- | --- | --- | --- | --- | --- | --- | --- |
|  |  |  | Normal | Low temp. | Out of range | Normal | Low temp. | Out of range | Normal | Low temp. | Out of range |  |  |
| Myanmar | Mar & Apr | 112529 | 66923 | 17641 | 27965 | 54476 | 18538 | 11550 | 43858 | 18472 | 10684 | 16569 | 45761 |
| Yunnan | Mar & Apr | 56120 | 6960 | 8871 | 40289 | 4592 | 7352 | 3884 | 3040 | 6655 | 2246 | 94 | 9601 |
|  | May | 47120 | 5963 | 1909 | 39248 | 3347 | 1514 | 3011 | 2265 | 1361 | 1235 | 28 | 3598 |
|  | Jun | 45296 | 5118 | 363 | 39815 | 3071 | 305 | 2086 | 2281 | 328 | 755 | 50 | 2559 |
|  | Jul | 45880 | 6818 | 268 | 38794 | 5136 | 131 | 1819 | 4186 | 89 | 992 | 31 | 4244 |
| Guizhou | Jun | 20400 | 8426 | 1062 | 10912 | 6445 | 1291 | 1752 | 5088 | 1246 | 1401 | 78 | 6256 |
|  | Jul | 21081 | 10254 | 268 | 10559 | 8685 | 218 | 1618 | 7615 | 219 | 1069 | 166 | 7668 |
| Thailand | Mar & Apr | 25925 | 14450 | 41 | 11434 | 9962 | 291 | 4238 | 6264 | 683 | 3306 | 472 | 6475 |
| Laos & Vietnam | Mar & Apr | 44225 | 15784 | 1655 | 26786 | 10060 | 2029 | 5350 | 6855 | 2617 | 2617 | 1064 | 8408 |
| Guangxi | Apr & May | 25620 | 10400 | 1926 | 13294 | 7277 | 2456 | 2593 | 5324 | 2846 | 1563 | 573 | 7597 |
| Guangdong | Apr & May | 24400 | 20397 | 893 | 3110 | 18312 | 1291 | 1687 | 15589 | 2012 | 2002 | 4353 | 13248 |
| Hunan & Jiangxi | May & Jun | 78915 | 61670 | 3406 | 13839 | 54783 | 3944 | 6349 | 48228 | 4653 | 5846 | 24608 | 28273 |
| Hubei & Anhui | May & Jun | 40222 | 25799 | 4796 | 9627 | 27369 | 6341 | 4080 | 23469 | 6340 | 3901 | 10033 | 19776 |
|  | Jul | 25575 | 19762 | 51 | 5762 | 17853 | 51 | 1909 | 15759 | 55 | 2090 | 2995 | 12819 |
| Jiangsu & Shandong | Jul | 18600 | 18258 | 59 | 283 | 16773 | 86 | 1458 | 14197 | 146 | 2516 | 4134 | 10209 |
| **Grand total** | | **624,933** | **290,071** | **43,175** | **291,687** | **241,663** | **45,795** | **52,960** | **198,460** | **47,660** | **41,322** | **65,248** | **186,492** |

**Supplementary Table S3.** Summary of the distance and direction of final endpoints from the origin for simulated trajectories.

| **Region** | **Periods** | **Height (m)** | **No. of endpoints** | **Distance ± standard error (km)** | **Rayleigh test** | | |
| --- | --- | --- | --- | --- | --- | --- | --- |
|  |  |  |  |  | **Direction (°)** | ***r*** | ***P*** |
| Myanmar | Mar & Apr | 500 | 1238 | 338±6 | 141 | 0.56 | <0.0001 |
| Myanmar | Mar & Apr | 750 | 2486 | 324±4 | 136 | 0.43 | <0.0001 |
| Myanmar | Mar & Apr | 1000 | 3321 | 306±4 | 119 | 0.34 | <0.0001 |
| Myanmar | Mar & Apr | 1250 | 4759 | 345±3 | 89 | 0.44 | <0.0001 |
| Myanmar | Mar & Apr | 1500 | 6250 | 392±3 | 74 | 0.58 | <0.0001 |
| Myanmar | Mar & Apr | 1750 | 7543 | 420±4 | 64 | 0.67 | <0.0001 |
| Myanmar | Mar & Apr | 2000 | 9330 | 413±4 | 59 | 0.76 | <0.0001 |
| Myanmar | Mar & Apr | 2250 | 10834 | 314±3 | 58 | 0.78 | <0.0001 |
| Yunnan | Mar & Apr | 500 | 140 | 432±16 | 203 | 0.94 | <0.0001 |
| Yunnan | Mar & Apr | 750 | 337 | 388±12 | 202 | 0.79 | <0.0001 |
| Yunnan | Mar & Apr | 1000 | 502 | 291±9 | 211 | 0.67 | <0.0001 |
| Yunnan | Mar & Apr | 1250 | 594 | 254±8 | 191 | 0.41 | <0.0001 |
| Yunnan | Mar & Apr | 1500 | 711 | 314±11 | 111 | 0.32 | <0.0001 |
| Yunnan | Mar & Apr | 1750 | 1146 | 380±13 | 64 | 0.6 | <0.0001 |
| Yunnan | Mar & Apr | 2000 | 2183 | 419±10 | 59 | 0.83 | <0.0001 |
| Yunnan | Mar & Apr | 2250 | 3988 | 364±7 | 58 | 0.93 | <0.0001 |
| Yunnan | May | 500 | 65 | 246±19 | 205 | 0.78 | <0.0001 |
| Yunnan | May | 750 | 136 | 197±13 | 208 | 0.53 | <0.0001 |
| Yunnan | May | 1000 | 144 | 155±9 | 229 | 0.25 | 0.0001 |
| Yunnan | May | 1250 | 156 | 170±11 | 126 | 0.08 | 0.3673 |
| Yunnan | May | 1500 | 143 | 305±18 | 66 | 0.41 | <0.0001 |
| Yunnan | May | 1750 | 313 | 531±23 | 55 | 0.52 | <0.0001 |
| Yunnan | May | 2000 | 708 | 719±17 | 56 | 0.72 | <0.0001 |
| Yunnan | May | 2250 | 1933 | 672±10 | 51 | 0.82 | <0.0001 |
| Yunnan | Jun | 500 | 7 | 284±73 | 211 | 0.68 | 0.0316 |
| Yunnan | Jun | 750 | 22 | 197±37 | 276 | 0.11 | 0.7626 |
| Yunnan | Jun | 1000 | 41 | 220±22 | 279 | 0.33 | 0.0114 |
| Yunnan | Jun | 1250 | 55 | 195±18 | 285 | 0.22 | 0.0716 |
| Yunnan | Jun | 1500 | 75 | 273±32 | 8 | 0.12 | 0.3217 |
| Yunnan | Jun | 1750 | 279 | 597±29 | 53 | 0.51 | <0.0001 |
| Yunnan | Jun | 2000 | 619 | 730±20 | 56 | 0.66 | <0.0001 |
| Yunnan | Jun | 2250 | 1461 | 709±12 | 54 | 0.76 | <0.0001 |
| Yunnan | Jul | 1000 | 8 | 253±58 | 41 | 0.93 | <0.0001 |
| Yunnan | Jul | 1250 | 48 | 510±51 | 30 | 0.84 | <0.0001 |
| Yunnan | Jul | 1500 | 209 | 614±31 | 23 | 0.64 | <0.0001 |
| Yunnan | Jul | 1750 | 773 | 677±18 | 32 | 0.59 | <0.0001 |
| Yunnan | Jul | 2000 | 1140 | 660±14 | 30 | 0.56 | <0.0001 |
| Yunnan | Jul | 2250 | 2066 | 601±9 | 28 | 0.53 | <0.0001 |
| Thailand | Mar & Apr | 500 | 49 | 166±16 | 89 | 0.28 | 0.0209 |
| Thailand | Mar & Apr | 750 | 151 | 234±11 | 351 | 0.35 | <0.0001 |
| Thailand | Mar & Apr | 1000 | 585 | 462±14 | 27 | 0.59 | <0.0001 |
| Thailand | Mar & Apr | 1250 | 2194 | 655±8 | 43 | 0.81 | <0.0001 |
| Thailand | Mar & Apr | 1500 | 3496 | 714±6 | 43 | 0.84 | <0.0001 |
| Laos & Vietnam | Mar & Apr | 500 | 143 | 252±12 | 283 | 0.82 | <0.0001 |
| Laos & Vietnam | Mar & Apr | 750 | 290 | 498±21 | 292 | 0.56 | <0.0001 |
| Laos & Vietnam | Mar & Apr | 1000 | 1087 | 656±13 | 354 | 0.53 | <0.0001 |
| Laos & Vietnam | Mar & Apr | 1250 | 2477 | 705±9 | 22 | 0.63 | <0.0001 |
| Laos & Vietnam | Mar & Apr | 1500 | 4411 | 756±7 | 33 | 0.73 | <0.0001 |
| Guangxi | April & May | 500 | 40 | 260±19 | 298 | 0.44 | 0.0003 |
| Guangxi | April & May | 750 | 368 | 570±21 | 14 | 0.6 | <0.0001 |
| Guangxi | April & May | 1000 | 1302 | 631±11 | 24 | 0.7 | <0.0001 |
| Guangxi | April & May | 1250 | 2395 | 636±8 | 23 | 0.7 | <0.0001 |
| Guangxi | April & May | 1500 | 3492 | 659±7 | 27 | 0.74 | <0.0001 |
| Guangdong | April & May | 500 | 1318 | 289±7 | 341 | 0.34 | <0.0001 |
| Guangdong | April & May | 750 | 2634 | 500±7 | 9 | 0.56 | <0.0001 |
| Guangdong | April & May | 1000 | 3401 | 530±6 | 16 | 0.59 | <0.0001 |
| Guangdong | April & May | 1250 | 3128 | 532±6 | 22 | 0.6 | <0.0001 |
| Guangdong | April & May | 1500 | 2767 | 521±7 | 33 | 0.63 | <0.0001 |
| Guizhou | Jun | 750 | 18 | 612±99 | 49 | 0.73 | <0.0001 |
| Guizhou | Jun | 1000 | 103 | 430±29 | 317 | 0.17 | 0.0473 |
| Guizhou | Jun | 1250 | 385 | 412±13 | 325 | 0.45 | <0.0001 |
| Guizhou | Jun | 1500 | 709 | 443±10 | 335 | 0.52 | <0.0001 |
| Guizhou | Jun | 1750 | 1224 | 455±8 | 336 | 0.49 | <0.0001 |
| Guizhou | Jun | 2000 | 1638 | 453±7 | 345 | 0.52 | <0.0001 |
| Guizhou | Jun | 2250 | 2179 | 425±6 | 349 | 0.49 | <0.0001 |
| Guizhou | Jul | 750 | 48 | 824±67 | 38 | 0.84 | <0.0001 |
| Guizhou | Jul | 1000 | 165 | 549±32 | 22 | 0.41 | <0.0001 |
| Guizhou | Jul | 1250 | 520 | 512±17 | 352 | 0.43 | <0.0001 |
| Guizhou | Jul | 1500 | 988 | 567±12 | 0 | 0.49 | <0.0001 |
| Guizhou | Jul | 1750 | 1635 | 635±10 | 2 | 0.49 | <0.0001 |
| Guizhou | Jul | 2000 | 1990 | 645±9 | 5 | 0.5 | <0.0001 |
| Guizhou | Jul | 2250 | 2322 | 605±8 | 3 | 0.47 | <0.0001 |
| Hunan & Jiangxi | May & Jun | 500 | 1632 | 452±7 | 348 | 0.5 | <0.0001 |
| Hunan & Jiangxi | May & Jun | 750 | 4527 | 482±5 | 336 | 0.39 | <0.0001 |
| Hunan & Jiangxi | May & Jun | 1000 | 6406 | 484±4 | 333 | 0.36 | <0.0001 |
| Hunan & Jiangxi | May & Jun | 1250 | 7638 | 469±3 | 340 | 0.36 | <0.0001 |
| Hunan & Jiangxi | May & Jun | 1500 | 8070 | 447±3 | 354 | 0.37 | <0.0001 |
| S Hubei & S Anhui | May & Jun | 500 | 1797 | 542±8 | 354 | 0.46 | <0.0001 |
| S Hubei & S Anhui | May & Jun | 750 | 3102 | 511±6 | 334 | 0.34 | <0.0001 |
| S Hubei & S Anhui | May & Jun | 1000 | 3891 | 471±5 | 318 | 0.28 | <0.0001 |
| S Hubei & S Anhui | May & Jun | 1250 | 4904 | 427±4 | 302 | 0.26 | <0.0001 |
| S Hubei & S Anhui | May & Jun | 1500 | 6082 | 382±4 | 296 | 0.27 | <0.0001 |
| Hubei & Anhui | Jul | 500 | 1654 | 667±12 | 6 | 0.67 | <0.0001 |
| Hubei & Anhui | Jul | 750 | 2168 | 670±10 | 359 | 0.51 | <0.0001 |
| Hubei & Anhui | Jul | 1000 | 2546 | 649±9 | 350 | 0.37 | <0.0001 |
| Hubei & Anhui | Jul | 1250 | 3028 | 623±8 | 330 | 0.27 | <0.0001 |
| Hubei & Anhui | Jul | 1500 | 3423 | 615±7 | 307 | 0.25 | <0.0001 |
| Jiangsu & Shandong | Jul | 500 | 1518 | 683±11 | 354 | 0.57 | <0.0001 |
| Jiangsu & Shandong | Jul | 750 | 1971 | 691±10 | 359 | 0.47 | <0.0001 |
| Jiangsu & Shandong | Jul | 1000 | 2137 | 679±10 | 349 | 0.31 | <0.0001 |
| Jiangsu & Shandong | Jul | 1250 | 2239 | 652±9 | 321 | 0.22 | <0.0001 |
| Jiangsu & Shandong | Jul | 1500 | 2344 | 645±8 | 301 | 0.2 | <0.0001 |

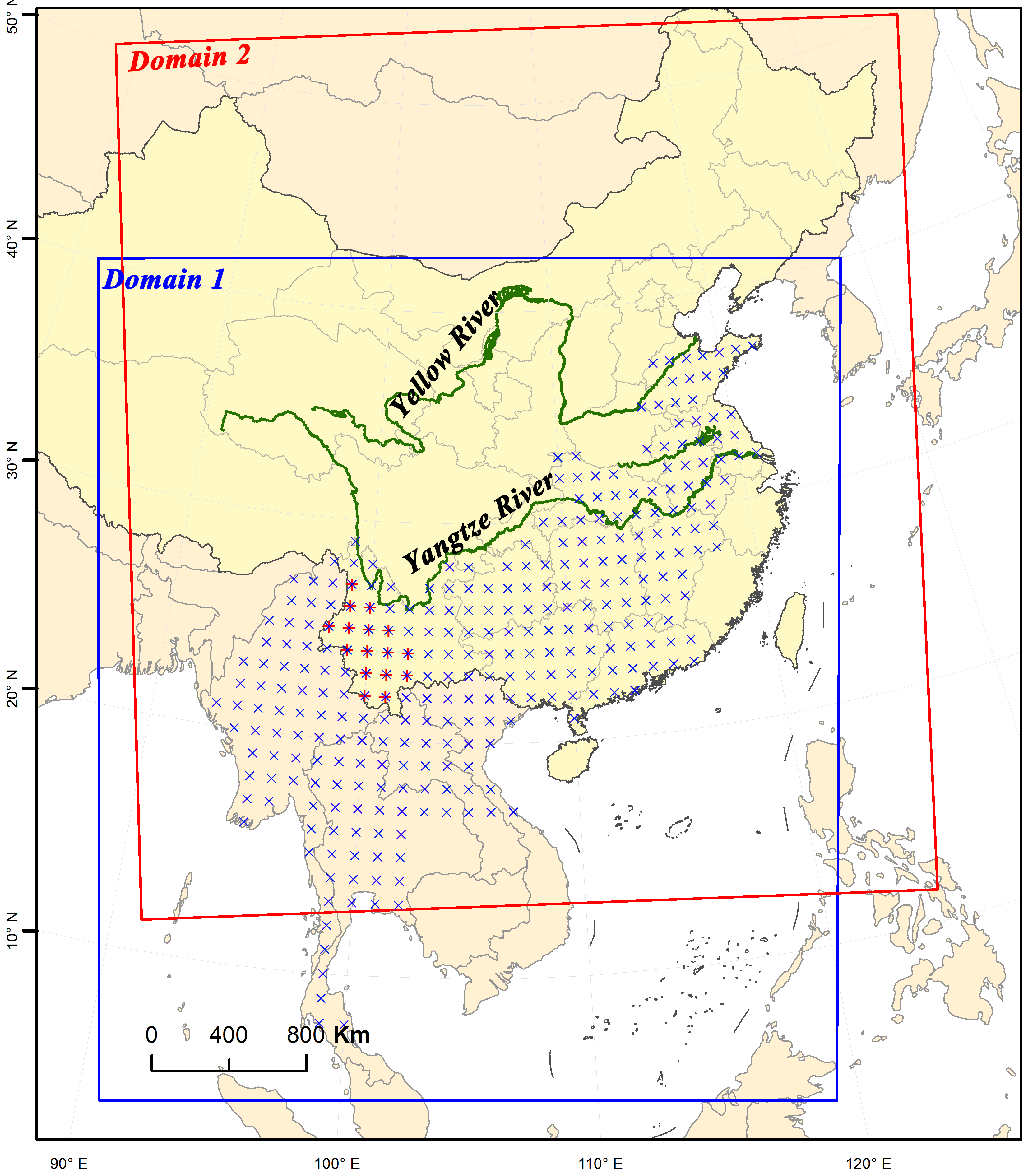

**Supplementary Figure S1.** Study area used in the WRF model (blue and red squares) for the trajectory analyses, and location of the origins of trajectory simulations (blue and red points). FAW found in Yunnan were restricted to the southwestern region in January–March 2019, thus the trajectories in that period were only started from southwestern Yunnan (red points). Domain 1 was used in the trajectory simulation for Indochina and Yunnan in March and April, and Domain 2 was used for China from April to July.
